# Spatial structure drives the evolution of parasites into host defense systems

**DOI:** 10.64898/2026.10.14.759074

**Authors:** Sultan Ahmed Nazir, Ben Ashby, Asher Leeks, Bram van Dijk

## Abstract

Host defense systems in prokaryotes and eukaryotes often appear to be derived from the very parasites they now defend against, yet the evolutionary route for such a transition remains poorly understood. Based on the biology of integrating viruses such as temperate bacteriophages and retroviruses, we develop a multilevel eco-evolutionary framework in which within-host viral replication is treated as a public goods game over a costly, shared replication product. In well-mixed host populations, we predict convergence on a single viral strategy of intermediate replicative investment. However, when we impose spatial structure by restricting host reproduction and virus transmission to local events, the virus population diversifies into two lineages: a horizontally transmitted lineage that parasitises hosts, and a vertically transmitted hyperparasitic lineage that parasitises the horizontally transmitting viruses. The latter “hyperparasite” is a viral cheat that suppresses the replication and propagation of the parasitic virus, thereby acting as a defense system to the hosts. Local interactions concentrate the benefits of this suppression among neighbouring hosts that carry related proviruses, allowing selection on the proviral cheat as a host-level defense. Our results show that evolutionary cheating within parasite populations can drive the evolution of parasites into defense systems in spatially structured host populations.

**SIGNIFICANCE STATEMENT:** Parasites are traditionally defined as evolutionary antagonists, but a growing body of evidence shows that parasitic elements can be repurposed into host defense systems. How these transitions happen is poorly understood. Using a model of social evolution of integrating viruses, we find selection for a vertically inherited lineage of viral cheats in spatially structured host populations. These integrated cheats, from the perspective of the host, act as an ‘enemy of my enemy’ by suppressing the transmission of free viruses. Our results provide novel insights into the outcomes of viral evolution and a theoretical mechanism for the origin of parasite-derived host immunity.

## INTRODUCTION

Parasitism occurs across the tree of life. Once parasites establish in a host population, infected hosts become a resource in their own right: other parasites can feed on nutrients leaking from damaged tissue or hijack the first parasite’s machinery. As a result, “parasites of parasites”, or hyperparasites, emerge as a predictable consequence of parasitism (Parratt & Laine, 2016, 2018; Poelman et al., 2022; Yau et al., 2011). At the same time, host lineages have evolved diverse defense mechanisms against their parasites, often undergoing rapid coevolution. In many cases, host defense mechanisms are phylogenetically derived from genes found in parasites themselves (Broecker, 2026). For example, in prokaryotes, many antiphage defense systems are carried by integrated viruses (prophages) and other mobile genetic elements (Getz & Maxwell, 2024; Koonin et al., 2020; Kuang et al., 2026), integrated virophages defend unicellular eukaryotes against giant viruses (Talha et al., 2026), and in vertebrates, many immune functions are derived from transposable elements and viral elements found within the genome (Broecker & Moelling, 2019; Grandi & Tramontano, 2018; Koonin & Krupovic, 2015). Defensive microbial symbionts of eukaryotes also commonly emerge from pathogens (Ashby & King, 2017; Drew et al., 2021; Sachs et al., 2011). Together, these examples reveal a recurring evolutionary pattern: evolution of defense systems from an *enemy of my enemy* dynamic.

To become a host defense system, a hyperparasite must be vertically inherited from the host to its offspring, for instance by integrating into the host genome. Indeed, many *enemy of my enemy* defense systems have evolved in the types of parasites that already undergo some vertical transmission: temperate phages can integrate into bacterial chromosomes as prophages, and are then inherited when the cell divides; retroviruses and transposable elements can enter the host germline in multicellular organisms (Boeke & Stoye, 1997; Tracy & Zhang, 2025); and plasmids and episomes can be vertically inherited without integrating into chromosomes (Serquiña & Ziegelbauer, 2017). Even at the larger scale of microbiomes, vertical transmission of prokaryotic and fungal symbionts is an important evolutionary driver of host-symbiont relationships (Bright & Bulgheresi, 2010; van Vliet & Doebeli, 2019).

The protection offered by the vertically inherited hyperparasite can benefit hosts directly, in which the resident element blocks infection by the parasite within the same host, as in superinfection exclusion by resident prophages (Bucher & Czyż, 2024) or colonisation resistance by defensive symbionts (Wang et al., 2023), and thereby protects the hosts from the harmful effects of the parasite. Because this benefit is realised immediately and entirely by the individual carrying the element, direct defense can be favoured by individual-level selection, i.e. a host with the element outcompetes one without it. In other cases, the element inhibits the replication machinery of the parasite population, reducing the parasite’s overall output but without improving the survival of its own host. Such indirect defense is common in phage-encoded abortive infection defense systems (Owen et al., 2021) and defense mediated by satellite viruses (Fillol-Salom et al., 2020). Because the host carrying such an element gains no direct advantage, evolution of these defenses is harder to explain.

Here, we study how cheating within the parasite population, combined with spatial structure among hosts, can drive the evolution of parasite-derived, indirect defense systems. We focus our analysis on viruses for three reasons. First, viral cheats, such as variants that exploit the replication machinery of co-infecting viruses, are known to arise readily during viral infections (Leeks et al., 2021). Second, many viruses can integrate into their host genome as proviruses, allowing for a mix of horizontal and vertical transmission. Third, as discussed above, proviruses frequently contribute to host defense against other viruses. Integrating viruses therefore make an ideal setting to ask when and how viral cheats (the hyperparasites) can evolve into host defenses. We develop a multilevel model of viral evolution in which viruses within a host play a public goods game over costly, shared replication products. We analyse this model framework at increasing levels of biological complexity. In well-mixed host populations, adaptive dynamics predicts evolution towards a single, intermediate level of replicative investment (cooperation), to the detriment of hosts, consistent with other models of social evolution of viruses (Chao & Elena, 2017; Frank, 1994). However, in a spatially structured individual-based model, from sufficient standing viral diversity, we observe selection for two distinct viral lineages: a non-integrated, parasitic lineage that is primarily transmitted horizontally, and an integrated, hyperparasitic lineage that is vertically inherited. The hyperparasite is a viral cheat that suppresses replication of the parasitic virus, thereby indirectly protecting neighboring hosts from infection. Finally, a more detailed model that tracks individual virus particles shows conditions of viral lifecycle under which this defense can also evolve *de novo* from a single ancestral parasite. Together, our results show that mixed transmission modes and spatial structure allow host immunity to evolve from the ecological diversity within parasite populations.

## RESULTS

### A multilevel model of coinfections of integrating viruses

To study the social evolution of these viruses, we model an evolving viral trait, α ∈ [0, 1] that measures a focal viral lineage’s production of a shared (*trans-acting*) protein required for successful infection, such as a replicase enzyme or capsid protein. We assume that this protein is costly for an individual virus to encode, but that it benefits all viruses coinfecting the same host cell, and hence acts as an evolutionary public good (Chao & Elena, 2017; Frank, 1994; Leeks et al., 2021). The model has two coupled levels: host-level epidemiology (Figure 1A) and within-host virus replication (Figure 1B).

**Figure 1:**
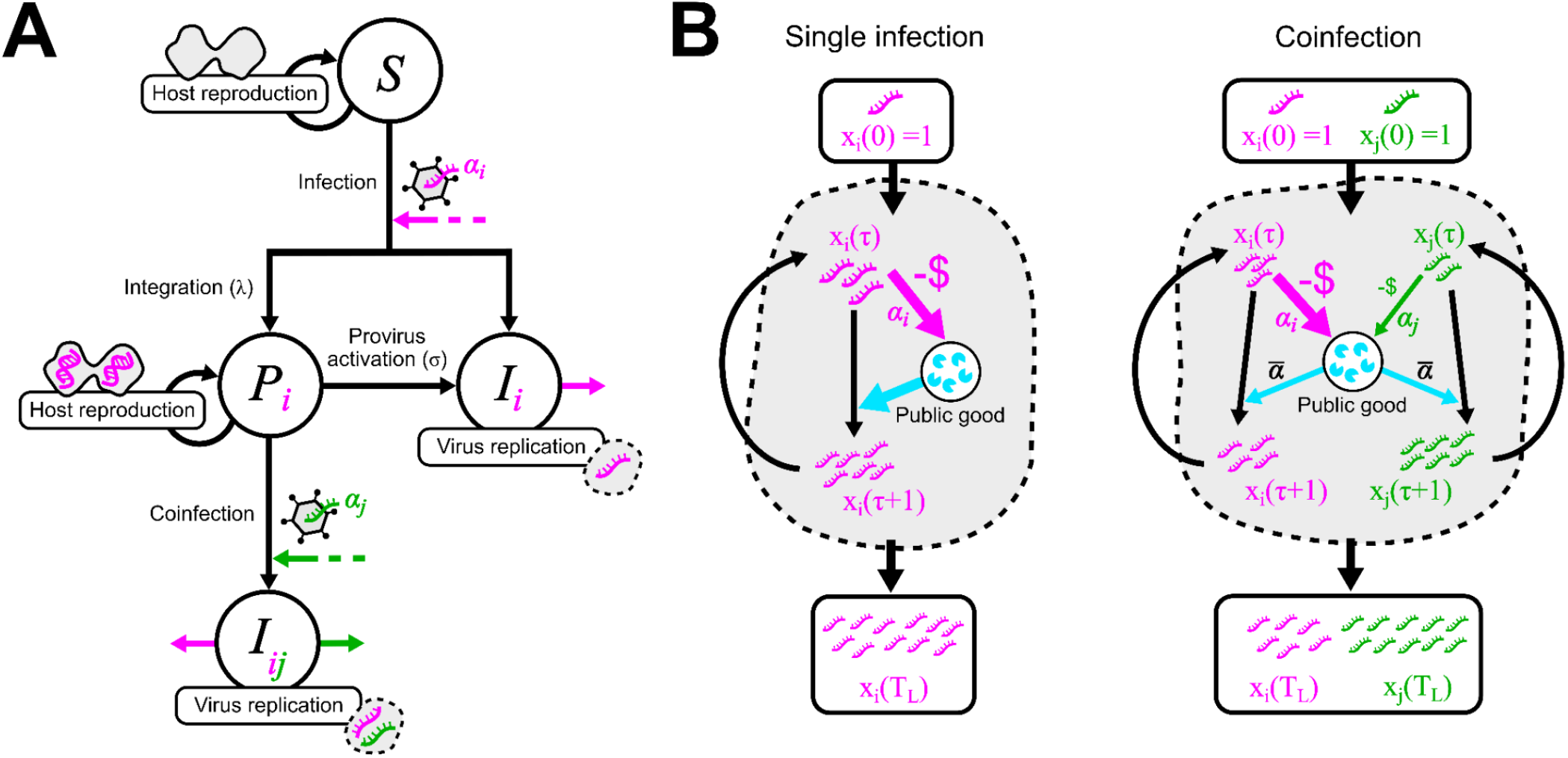
Overview of the eco-evolutionary model of integrating viruses. (A) Host level dynamics are described by a compartment model, and (B) the replication dynamics of viruses within a host are given by a public goods model. The total viral output at the end of the replicating period *T*_*L*_ determines the transmission rate of the virus in the host-level compartment model. All equations are given in the main text.

#### Host-level dynamics

Host-level dynamics follow a modified version of the coinfection model of van Baalen & Sabelis (1995), in which hosts can be uninfected, singly infected, or doubly infected. We extend this model to allow singly infected hosts to carry either a quiescent provirus or an actively replicating free virus. Uninfected, susceptible hosts occupy the *S* compartment. When a susceptible host is infected by a virus with trait value α_*i*_, the virus integrates to form a provirus with probability λ, moving the host to compartment *P*_*i*_; otherwise, the single virus begins replicating immediately, and the host moves to compartment *I*_*i*_. All hosts that are either uninfected or carry a quiescent provirus (i.e., in *S* or any *P*_*i*_) reproduce logistically with intrinsic birth rate *b* and die at intrinsic rate *d*. A provirus becomes active in two ways: spontaneously, at rate σ, or whenever a coinfection occurs. In the latter case, to ensure that a provirus interacts with the secondary infection, a host in *P*_*i*_ that is infected by a virus with trait value α_*j*_ has its resident provirus immediately activated. Both viruses are then replicating, and the host moves to the doubly infected compartment *I*_*ij*_. Once a host carries any actively replicating virus (i.e., is in *I*_*i*_ or *I*_*ij*_), it stops reproducing and dies at a rate γ > *d*. We denote the transmission rate of virus *α*_*i*_ from a singly infected host *I*_*i*_ as β_*i*_, and the transmission rates of viruses α_*i*_ and α_*j*_ from a doubly infected host *I*_*ij*_ as β_*ij*_ and β_*ji*_ respectively. Refer to Materials and Methods for the equations.

#### Within-host dynamics

Every viral copy within a host draws replication resources from a shared pool, to which each resident virus contributes in proportion to its cooperation trait α; each virus thus receives resources equal to the mean α within the host. Higher cooperation also entails a direct replication cost, which could reflect a trade-off between replication and transcription (Chao & Elena, 2017). If *x*_*i*_(τ) denotes the within-host abundance of a virus of trait value α_*i*_ at time step τ, then

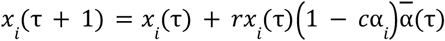

Where *r* is the maximum replication rate, *c* is the cost of cooperation, and 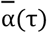 is the abundance-weighted mean trait value within the host. For singly-infected hosts in compartment 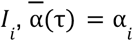, and for co-infected hosts in compartment 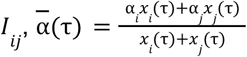.

We begin the replication dynamics with exactly one copy of every infecting virus (*x*_*i*_ (0) = 1 for hosts in *I*_*i*_ and *x*_*i*_ (0) = *x*_*j*_ (0) = 1 for hosts in *I*_*ij*_).

#### Coupling the two levels

We link the within-host and host-level models through the period *T*_*L*_, the duration of active viral replication before the host lyses. The host-level lysis rate is then γ = θ/*T*_*L*_, where θ is a conversion factor that translates within-host time steps into host-level time units. Because *T*_*L*_ is fixed across all infections, variation in the evolving trait α does not alter the timing of lysis; instead, it influences host-level dynamics solely through its effect on burst size, and hence transmission rate.

For simplicity, we approximate the transmission dynamics of viruses by setting their transmission rates proportional to the product of their burst size at internal time τ = *T*_*L*_ and the lysis rate γ. Thus, for a host in class *I*_*i*_, β_*i*_ = β_0_ γ_*i*_ *x* (*T*_*L*_) where *x*_*i*_ (*T*_*L*_) is the within-host abundance of singly infecting strain *i* at the end of the replication period. For a coinfected host in class *I*_*ij*_, β_*ij*_ = β_0_ γ_*i*_ *x* (*T*_*L*_) and β_*ji*_ = β_0_γ*x*_*j*_ (*T*_*L*_), where *x*_*i*_ (*T*_*L*_) and *x*_*j*_ (*T*_*L*_) denote the within-host abundances of coinfecting strains *i* and *j* at the end of the shared replication period.

See Supplementary Information S1 for the table summarising all model parameters.

### Spatially unstructured host populations lead to intermediate levels of viral cooperation

Using mutant invasion analysis, we found that under our standard set of assumptions, α evolved towards a single uninvadable strategy α_*opt*_, which satisfies

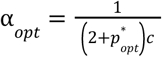

where *c* is the cost of cooperation. 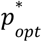 denotes the equilibrium fraction of susceptible hosts carrying a provirus of trait value α_*opt*_ under monomorphic viral dynamics (i.e., when all viruses share the same trait value). See Supplementary Information S2 for the complete derivation.

This intermediate optimum arises because two opposing forces act on viral cooperation (α). Within a coinfected host, viruses with lower α replicate faster than their more cooperative co-infecting partners, and so are transmitted preferentially, favouring ever-lower α. But across hosts, low-α viruses generate a smaller shared resource pool, and therefore produce a smaller total viral load, than more cooperative viruses. In other words, while selection at the level of viruses (within hosts) promotes the emergence of hyperparasites, higher-level selection of viral populations (between hosts) counteracts this to promote viral cooperation. In our model, increasing α beyond some point yields diminishing returns in viral output, so higher cooperation does not translate indefinitely into more viral copies. The relative strengths of these within-host and between-host forces influence the final level of cooperation that evolves.

#### Why the mean-field model favours a single optimum

When a virus exists as a provirus, its fitness is tied to its host’s reproductive fitness. The total fitness of a viral lineage combines vertical transmission (via host reproduction) and horizontal transmission (via infection of new hosts after within-host replication). Because the viral trait α does not affect the integration rate, induction rate, host death rate, or host reproduction rate, selection on α acts solely through horizontal transmission success. As also shown analytically in Supplementary Information S2, the adaptive dynamics optimum therefore corresponds to the α value that maximises horizontal transmission.

Pairwise invasibility plots confirm that the population evolves towards this optimum and that it is evolutionarily stable (See Figure 2). Note that this equation is not a closed expression for α_*opt*_. However, it reveals a quantitative relationship that the evolutionarily stable level of cooperation, α_*opt*_, increases as the corresponding equilibrium fraction of provirus carriers in the host population 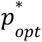 decreases. Because coinfection in our model can only occur when a virus infects a provirus-carrying susceptible host, *p* effectively measures the chance for coinfection, and so this relationship means that cooperation is favoured most strongly when coinfection is rare. Within the derivation for the optimum (See Supplementary Information S2), we observed that positive selection on α requires

**Figure 2:**
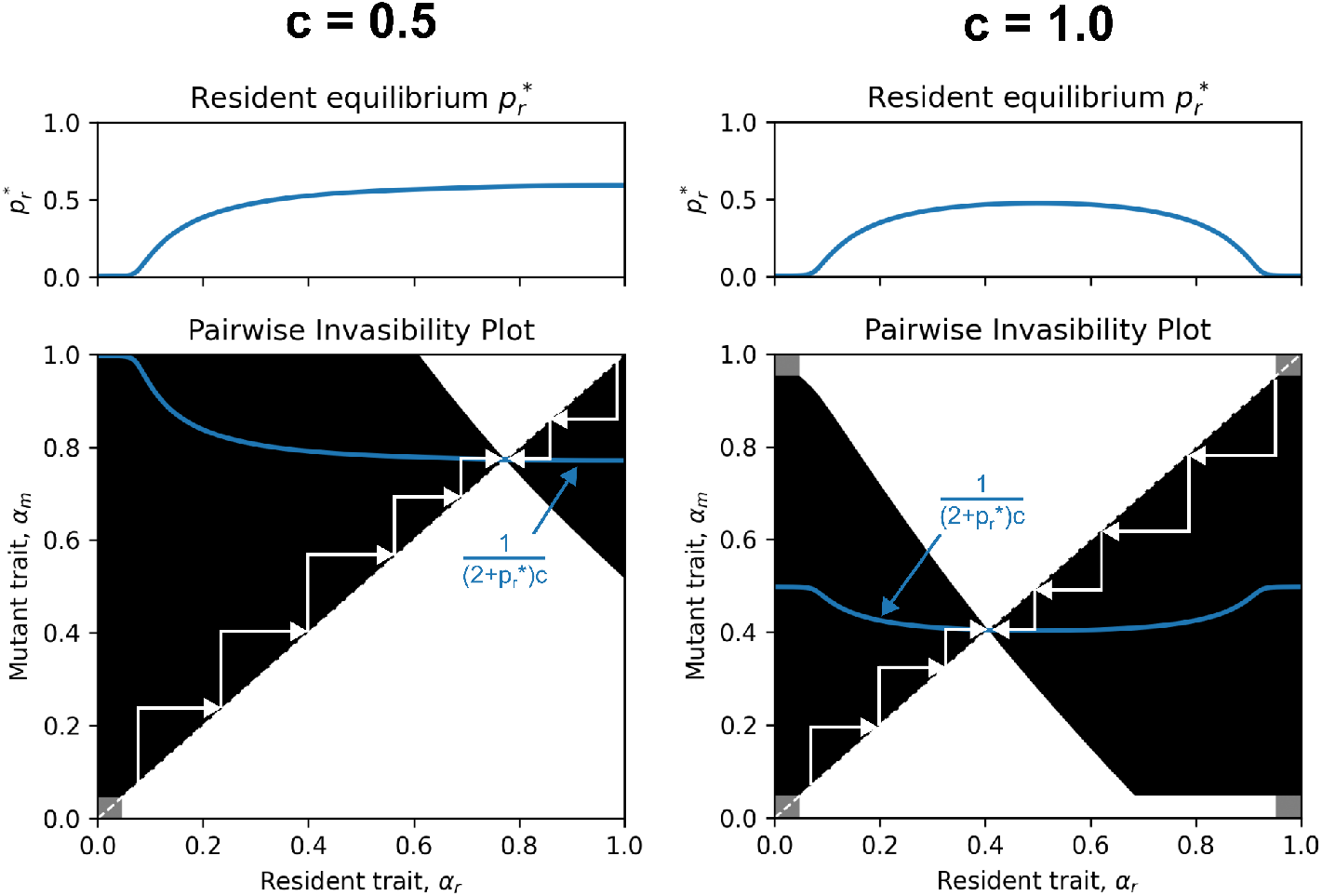
Pairwise invasibility plots. For two different values of cost of cooperation, *c* = 0. 5 and *c* = 1. 0, the equilibrium fraction of provirus-carrying susceptible hosts for a single resident virus is shown in the top row as a function of the virus trait value α_*r*_. The bottom row shows the pairwise invasibility plot showing the success of invasion by a mutant virus of trait value α_*m*_ against a resident virus of trait value α_*r*_. Black and white regions denote positive and negative invasion fitness respectively. The region is shaded in grey if both the resident and the mutant viruses have *R*_0_ < 1, in which case neither virus can persist. We observe that the virus evolves towards evolutionarily stable trait value that satisfies 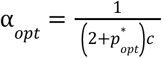

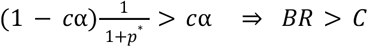

Where 1 – *c*α is the marginal benefit of cooperation (*B*) and *c*α is the marginal cost (*C*). The lineage-weighted relatedness at equilibrium (*R*) is given by 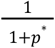 *R* approaches 0.5 if most infections are coinfections and 1 if coinfections are rare. As shown for public goods models (Frank, 2010), our result can therefore be thought of as a special case of Hamilton’s rule (Hamilton, 1964): coinfection lowers the relatedness between the viral variants sharing a host, since the benefits of cooperation are then split with an unrelated co-infecting virus rather than reaped exclusively by clonal copies of the cooperator, and thus lower relatedness weakens selection for cooperation.

### Host-level spatial structure promotes vertically transmitted viral cheats

In natural environments, bacteria often form spatially structured communities such as biofilms or soil microaggregates, where physical constraints and limited diffusion restrict both host reproduction and viral transmission to localised regions. To that end, we next simulated the same dynamics with an individual-based model on a square lattice. We initialised every lattice site with a susceptible host, half of which carried a provirus with a cooperation level (α) drawn uniformly between 0 and 1. Starting from this standing variation, we track which viral strategies persisted over time.

First, we confirmed that the lattice model preserves the mean-field behaviour of the earlier adaptive dynamics framework by assuming that host reproduction and virus transmission are global events. Despite effects of stochasticity, viral populations in this mixed regime indeed converged on the analytically derived optimum as a function of the cost of cooperation, *c* (Figure 3A). We then relaxed the mean-field assumption of global mixing by restricting host reproduction and virus transmission to local events among immediate neighbors on the lattice. This change produced a qualitatively different outcome. Instead of settling on a single viral strategy, two distinct viral strategies persist (Figure 3B and Figure S1): cooperators that are predominantly free viruses (shown in red), and cheats that are largely seen in a proviral state (shown in blue). This polymorphism is most clearly visible for intermediate values of the cost of cooperation (*c*).

**Figure 3:**
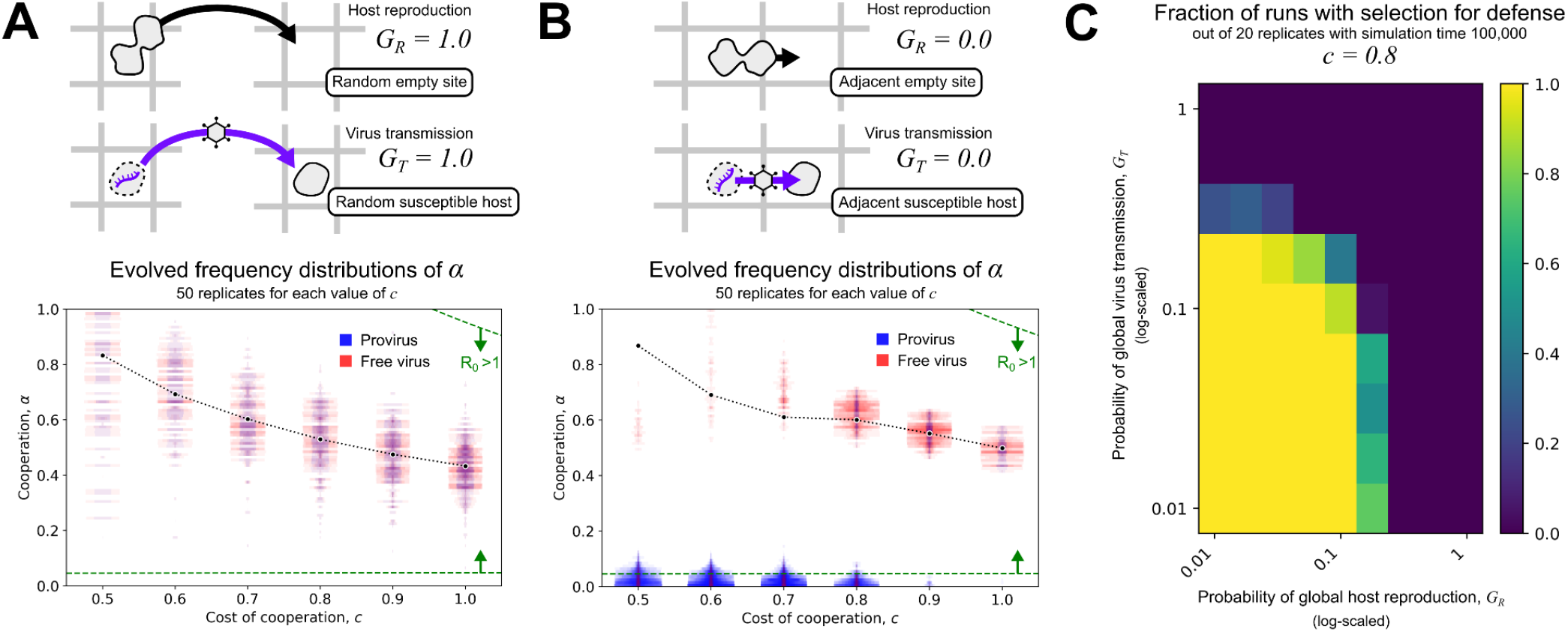
Local interactions result in polymorphism of two viral lineages. (A-B) After initialising the population with viruses of all types, we analysed the frequency distribution of viral types after 100,000 time steps under mixed (A) and spatial (B) conditions. The bottom distributions show the average abundance of α values within bins of size 0.05 from time steps 90,000 to 100,000, normalised by the highest abundance. Every singly infected host is counted once and every doubly infected host is counted twice for the α abundances. These histograms are visualised for different values of cost of cooperation, *c*. For each value of *c*, overlaid distributions of α values are shown for 50 different simulation runs. We excluded simulation runs where the virus went extinct before timestep 100,000. This includes 5 runs of *c* = 0. 5 in (A), and 3 runs of *c* = 0. 5 and 5 runs of *c* = 1. 0 in (B). For each simulation run without extinction, the peak-normalised distributions for viruses found in ‘provirus’ and ‘free virus’ states, are plotted in blue and red respectively, which appear purple when they overlap. Each black dot is the average predicted α_*opt*_ over the replicates. We considered the fraction of provirus-carrying susceptible hosts averaged over time steps 90,000 to 100,000 to be equal to 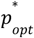 to estimate α_*opt*_ for each replicate. The green dotted line represents the α value at which *R*_o_ of the virus is equal to one. See Supplementary Information S3 for the derivation of *R*_0_. (C) Heatmap of the fraction of simulation runs, out of 20 replicates, where the group of proviral cheats (i.e. defense system) emerged for different probabilities of global host reproduction (*G*_*R*_) and global virus transmission (*G*_*T*_).

### Why spatial structure results in a survival of two lineages

The results above showed that local host reproduction and local virus transmission resulted in the emergence of two lineages, rather than one. When a low-alpha provirus (a viral cheat) is activated by an incoming free virus, it may reduce the viral output during within-host replication, but the host is bound to die either way. However, spatial structure results in an additional relatedness structure among hosts (Best et al., 2010; Boots & Sasaki, 1999). The reduced viral output is therefore protecting related individuals nearby, hosts that are likely to also carry the vertically transmitted viral cheat. In other words, the proviral cheat acts as an indirect defense system: it is favoured not through its own horizontal transmission, but through the protection it offers to groups of related hosts, and thereby to copies of itself. Neither condition suffices alone (Figure 3C): without local reproduction the neighbors are unrelated, and without local transmission the protection is spread across the whole population. In the mixed (mean-field) system, this structure disappears entirely, and with it any selection for proviral cheats.

To demonstrate this mechanism, we initialised simulations with patches of hosts carrying proviruses of different α values, and exposed them to an incoming wave of free viruses (Figure 4). Indeed, only non-cooperating proviruses (low α) were able to survive. While hosts with integrated viral cheats inhibit the transmission of free viruses, the converse can be said about hosts carrying integrated viral cooperators. Activated proviruses that cooperate (high α) enhance the replication and transmission of free virus, which preferentially kills hosts carrying the same provirus. Consequently, the cooperating viruses may occasionally integrate and form a provirus, but this vertically transmitted state cannot persist for long, explaining why they are found predominantly as free viruses (Figure 3B). In other words, the two viral types differ not only in α, but in their life cycle, and represent functionally distinct lineages. In this context, the labels that we assign to these viruses are dependent on the level of interaction: what is a *cooperator* to a coinfecting virus is a *parasite* to the host, and what is a *cheater* to the coinfecting virus is a *defense system* to the host.

**Figure 4:**
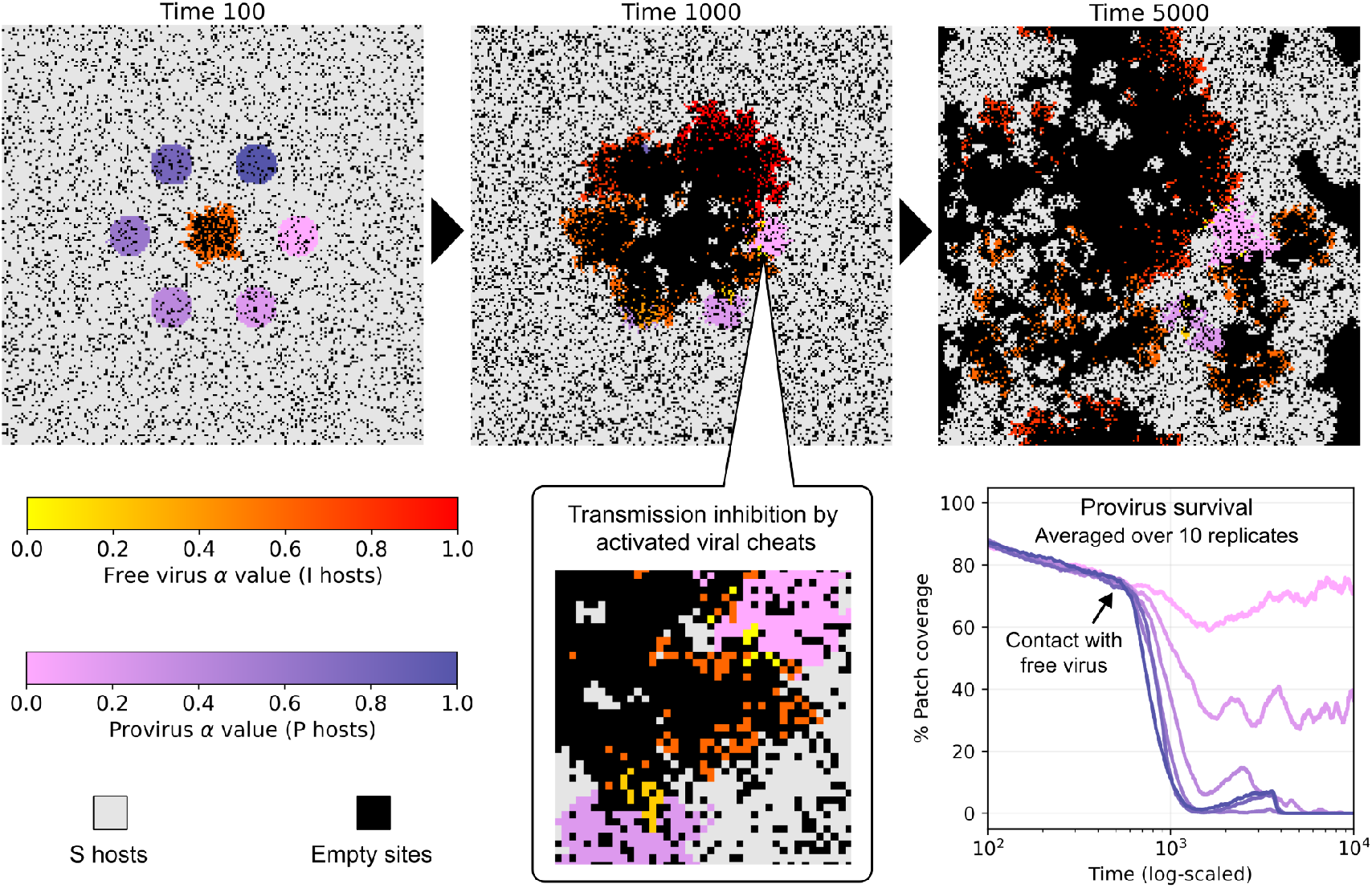
Demonstration of the selection for low-α provirus. Simulation experiment showing the survival of patches of proviruses of different α values when faced with invasion from a high *R*_0_ free virus. For the purpose of this demonstration, we initialised the simulation with equally sized circular patches of hosts carrying proviruses, each with a different value of α, at equal distances from the free virus outbreak. We observe that the patches of high-α proviruses amplify viral output, which wipes out the neighbouring provirus-harbouring hosts and speeds up the wave of free virus. On the other hand, low-α proviruses act like defense systems by inhibiting the replication of free viruses by the tragedy of the commons (i.e. exploitation of public goods by the cheaters drives the free virus population to extinction locally). Hence, these proviruses protect neighboring provirus-harbouring hosts and inhibit the transmission of the wave of free virus. In the long run, host-dense regions emerge that are rich in low-proviruses. The lattice point is coloured by the α value of its provirus for every host in state *P*, and by the α value of the free virus that initiated virus replication within hosts in state *I*. (Bottom-right) Average coverage of the patches with hosts carrying the assigned provirus as a function of time. For this experiment, we disabled any mutation, integration or spontaneous provirus activation, and *c* = 0. 5.

The lattice model demonstrated the establishment of a three-player ecological community: hosts without defense (S), hosts with defense (P), and free parasites (I). We have shown that hosts with defense (P) suppress transmission of the free parasite (I), while the free parasite (I) kills hosts without defense (S) to produce more free parasites (I). Although coexistence among players linked by such antagonistic interactions is inherently fragile and sensitive to the relative strengths of these interactions, spatial structure promotes their coexistence by connecting subpopulations with out-of-phase population cycles (Briggs & Hoopes, 2004; van Dijk et al., 2021).

### *De novo* evolution of parasite-derived defense system

The lattice model demonstrated that spatial structure provides a novel niche: proviral cheats that act as a host defense system. However, this model initialised with all possible viral variants present from the beginning, and therefore cannot explain the *de novo* evolution of such proviral cheats. Interestingly, when including mutations in the lattice model, we found that proviral cheats did not emerge from a single ancestral parasite, except when mutation rates were exceptionally high (see Figure S2). In addition, one may wonder if the simplification of within-host dynamics directly determining transmission rates is biologically relevant.

To better understand the *de novo* evolution of proviral cheats, and to further relax several simplifying assumptions made in the lattice model, we extended our model with more biological detail in virus-host interactions. Rather than approximating viral dynamics implicitly through transmission rates, our *virus factory model* explicitly tracks individual particles as they transition between three states: integrated provirus, actively replicating virus within a “virus factory”, and free extracellular virion (Figure 5(A)). A virus factory here refers to the pool of co-replicating viruses within a host, whose replication probabilities are shaped by the same public goods dynamics described earlier (see Methods and Supplementary Information S6 for more details).

**Figure 5:**
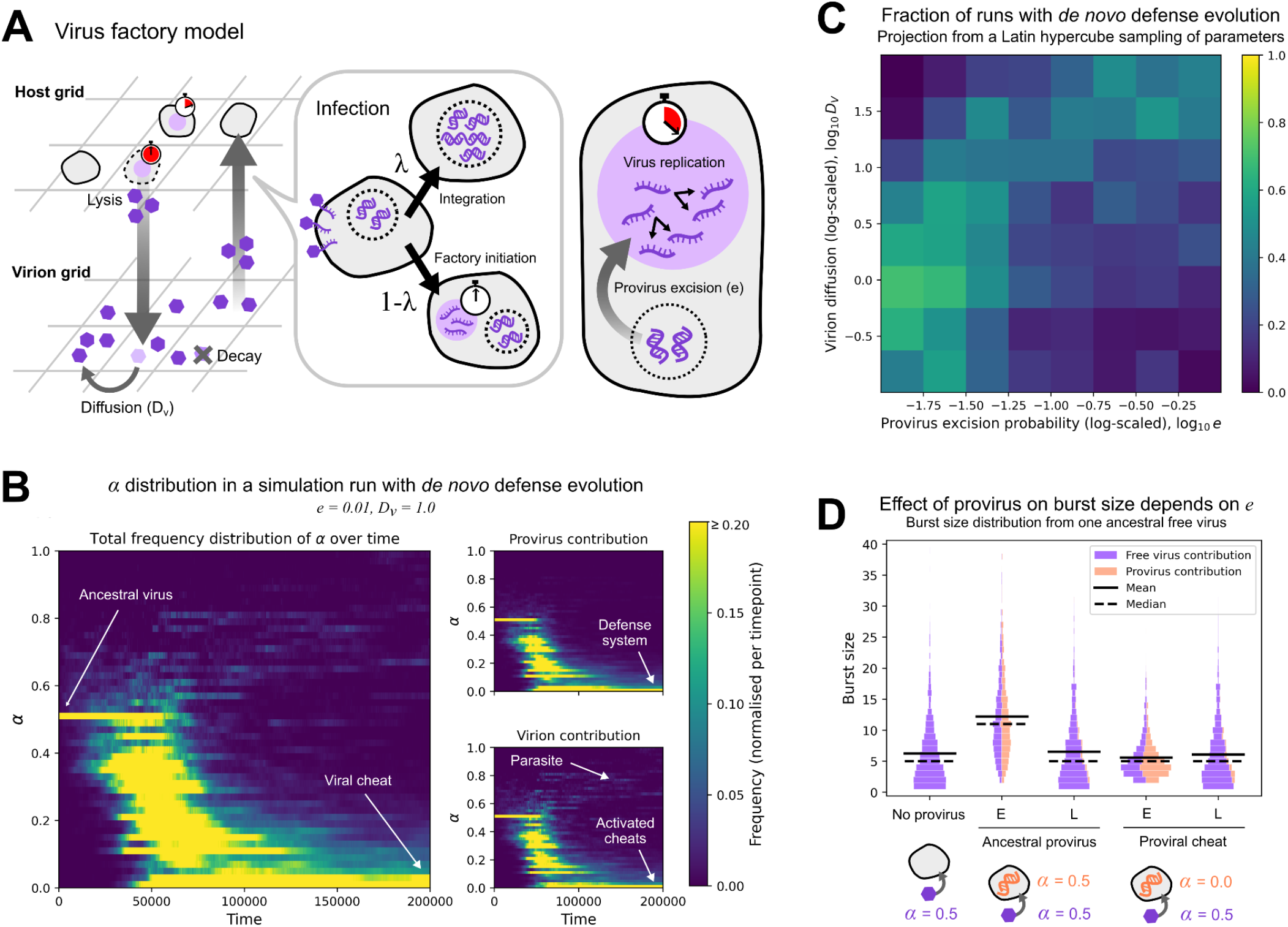
Virus factory model and *de novo* evolution of defense system. (A) Overview of the virus factory model. (B) One simulation run showing the evolution of normalised α frequency distribution where defense evolved *de novo*. (C) A heatmap of the projection from a Latin hypercube sampling of model parameters onto the parameters *D*_*V*_ and *e*, with each bin coloured by the fraction of simulation runs that fall within the bin that showed the evolution of viral cheats, when the virus population was initialised as a monomorphic population of α = 0. 5. See Supplementary Information S4 for sensitivity analysis over the other parameters. See Methods for a full description of the analysis. (D) Distribution of the burst size from 1000 replicates of virus factories all of which were initialised by one free virus of α = 0. 5. The distribution varies depending on the presence of a provirus in the host, its excision probability and the trait value of the provirus. E = early excision (*e* = 1) and L = late excision (*e* = 0. 01). Early excision increases the burst size if the provirus is also of the ancestral form. If the provirus is a viral cheat (α = 0), early excision suppresses the proportion of parasites. The bar corresponding to each burst size is divided by the average contribution of the free virus (purple) and the activated provirus (orange).

**Figure 6.**
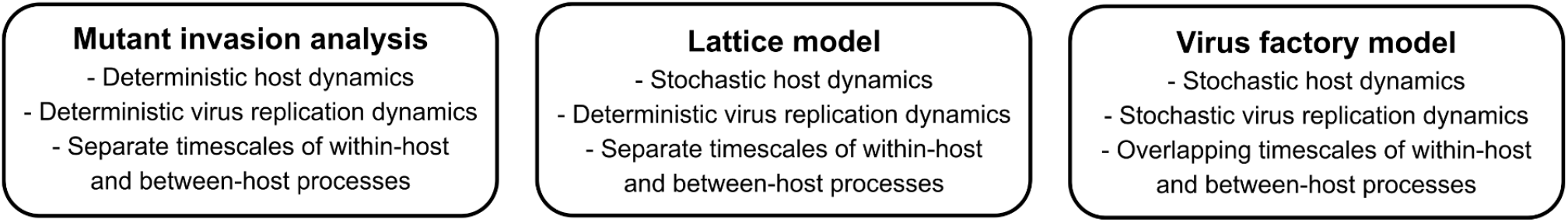
Modelling formalisms for host and virus dynamics used in this study.

Because the virus factory model has many more parameters, we first explored its behaviour broadly, varying six key parameters simultaneously with Latin hypercube sampling (Supplementary Information S7). These covered the provirus life cycle (integration, spontaneous induction, and the probability of joining an existing virus factory) and the extracellular environment (virion entry, diffusion and decay). Defensive proviral cheats emerged across a wide range of these parameter combinations, but, as in the lattice model, never when the system was well mixed. Over long evolutionary timescales, the system often settled into a state with very few high-α viruses remaining (see Figure 5B), suggesting that the evolved defense is highly effective at suppressing the parasitic virus.

Our parameter sweep also revealed a key obstacle to the emergence of defense: its evolution depends strongly on how early a provirus joins an active virus factory, set by the excision probability *e*, in combination with virion diffusion (Figure 5C). Since we assume the factory has a fixed lifetime, a provirus lineage that joins earlier participates in more rounds of replication and thereby exerts a stronger influence on the burst size and its composition. When a free virus initiates a virus factory, early excision of the provirus has opposing effects on the burst size depending on the trait value of the provirus (See Figure 5(D)). To understand this, consider what happens upon infection by the ancestral (cooperative) virus. If the provirus is also cooperative, early excision increases the abundance of the parasite because both produce public goods that contribute to the viral factory. If the provirus is instead a viral cheat, early excision reduces the abundance of the parasite, because the viral cheat is replicated without contributing to the viral factory.

With limited virion diffusion *D*_*V*_, both effects (extreme burst sizes due to cooperative proviruses, and reduced burst size due to cheating proviruses) fall on neighboring hosts carrying the same provirus. Therefore, early provirus excision has two opposing effects on the *de novo* evolution of defense: it strengthens selection on a proviral cheat, but imposes a cost in the maintenance of a cooperative proviral lineage. This cost is most relevant precisely when asking how defense can evolve *de novo*, since that scenario necessarily starts from a viral population without a defensive viral cheat.

Finally, the virus factory model relaxes two further assumptions of our earlier framework. First, within-host and between-host timescales are no longer separated: within-host replication still follows the public-goods dynamics described earlier, but at the end of the replication period the host lyses, releasing its viral particles into an overlying lattice of extracellular virions that diffuse and decay, with infection occurring when susceptible hosts pick up free virus. Second, hosts can now be infected by more than one free virus at a time and can carry more than one provirus. In other words, beyond showing the conditions for *de novo* evolution of the defense system, our virus factory model demonstrates that the mechanism of evolution of defensive proviral cheats is robust to the simplifying assumptions of the earlier model.

## DISCUSSION

In this study, we show that vertically transmitted viruses can evolve into host defense systems. Through a public goods game between viral lineages, vertically transmitted viral cheats can emerge that suppress the horizontal transmission of free viruses. Since these proviral cheats cannot persist through horizontal transmission, their fitness is tied to the survival and reproduction of their hosts. Therefore, parasite suppression in spatially assorted hosts leads to preferential protection of other proviral cheats. Using a mixture of analytical, numerical, and agent-based simulation models, we show that host-level spatial structure is required for the evolution and maintenance of these defenses. Our most biologically detailed model (the “virus factory model”) revealed an additional factor: the precise moment when a provirus begins exploiting the incoming virus during coinfection. Exploiting earlier gives stronger parasite suppression to protect neighboring hosts, but the same early-acting strategy accelerates parasite transmission for an ancestral parasitic provirus. In other words, for a fixed excision rate, the life-history conditions that make an established provirus a more effective defense can impede the initial evolution of defense from an ancestral parasite. Overall, host-level spatial structure and a viral public goods game, two conditions likely widespread in nature, consistently favour selection for defensive proviruses. Whether this selection drives a parasite-hyperparasite branching, however, depends on the stability of the proviral lineages. Therefore, we describe a general route by which parasite-derived elements can evolve into host defenses.

Our model is most relevant to the evolution of defense systems whose defensive function is a direct byproduct of their own replication. For example, virophages in eukaryotic cells are known to hijack the replication machinery of giant viruses within their virus factories (Talha et al., 2026), while phage satellites such as phage-inducible chromosomal islands (PICIs) in bacteria interfere with viral replication to function as innate immune systems (Dearborn et al., 2017; Fillol-Salom et al., 2020; Hays & Seed, 2020; Quiles-Puchalt et al., 2014). Crucially, virophages and PICIs are vertically transmitted – they reside in the host genome until another parasite induces them. The results from our model therefore show how such defensive viruses or sub-viral agents could have evolved from fully functional parasitic viruses. The relatively simple conditions required for our mechanism to work, such as mixed-mode viral transmission, viral public goods, and host spatial structure, could therefore provide one explanation for why so many defense systems have replication tied to the defensive phenotype.

In other host defense systems, the genes that confer defenses are not themselves replicators competing with the incoming virus. CRISPR-Cas systems, for example, also use integrated viral sequences to express a defensive phenotype, but the spacers they carry are not replicators that compete with the incoming virus, and the system gains no preferential amplification from defending against it (Barrangou, 2013). One possibility is that our mechanism drove the original evolution of such systems, and that the link between replication and defense was lost as the elements became further domesticated by the host. Alternatively, these systems may have a different origin altogether. In particular, defenses that directly protect the cell carrying them can be selected at the level of the individual host and do not depend on the spatial assortment that our mechanism requires. Indeed, modeling defense as an evolving host trait, Iranzo et al. (2015) showed that such immunity can evolve even in well-mixed populations, whereas altruistic defenses require spatial structure. Our results build on this by showing that spatial structure also enables altruistic defense systems to emerge not from *host-level evolution*, but from *parasite-level evolution*.

Our model focused on the origins of defense systems from parasites, but did not explore the longer-term eco-evolutionary consequences. For example, once a parasite-derived defense system is established, it becomes part of the selective environment of the parasite itself. Hyperparasitism could then shape selection on the parasite’s transmission rates. This co-evolution could yield interesting ecological dependencies, with parasites evolving extremely high levels of transmission or virulence that are only sustainable in the presence of a hyperparasite (Nowak & May, 1994; Smith & Ashby, 2023; Wood & Ashby, 2023). Another interesting co-evolutionary player to explore in future studies is the host, which we purposefully left out of our study. The host could for example evolve to domesticate the proviral cheat to avoid reversion to parasitism. Future models could investigate the outcomes of coevolution of the viral lifecycle traits like integration and provirus activation, along with its evolution on the parasite-mutualist spectrum, including whether these can give rise to defense systems in which replication and defensive phenotypes are no longer tied together. As shown in the context of the influence of provirus excision rate on *de novo* evolution of defense, different modes of transmission impose significant ecological constraints that influence the subsequent evolutionary trajectories. Taken together, we suggest that longer-term coevolutionary dynamics following from the mechanism we model could help to explain the diversity of parasite-derived defense systems.

Social evolution theory is increasingly being applied to viral populations (Díaz-Muñoz et al., 2017; Leeks et al., 2023). Our model extends previous work by including both a viral public goods game within cells and a vertical transmission component. As a result, we explore social consequences not just among viruses coinfecting a cell, but also between viruses integrated inside the genomes of neighboring host cells. This additional component reveals that there are two components of fitness that can act on viral cheats: direct fitness benefits to the cheating lineage, by increasing its competitiveness when coinfected by a cooperative virus; and indirect fitness benefits, by protecting related proviruses integrated inside neighboring host cells. In our model incorporating the possibility for indirect fitness benefits of cheating via spatial structure, viruses shifted from a mixed strategy of partial cooperation towards two distinct strategies: horizontally transmitted highly virulent cooperative viruses, and vertically transmitted low-virulence full cheats which resembled host defense systems. Our results therefore highlight the potential explanatory power from incorporating viral social interactions that occur over multiple spatial scales.

## MATERIALS AND METHODS

### Mutant invasion analysis

#### Resident population dynamics

For a resident virus with trait value α_*r*_,

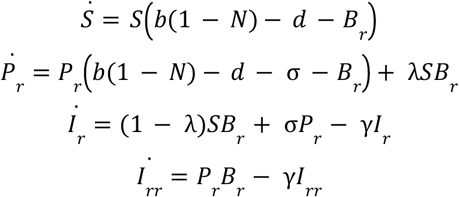

where *B*_*r*_ = β_*r*_ *I*_*r*_ + 2β_*rr*_ *I*_*rr*_ and *N* = *S* + *P*_*r*_ + *I*_*r*_ + *I*_*rr*_.

#### Mutant population dynamics

Assuming the resident population dynamics has reached equilibrium with steady state abundances 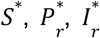 and 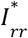 such that 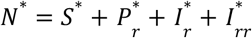 and 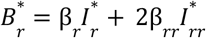, then, for a rare mutant virus with trait value α_*m*_,

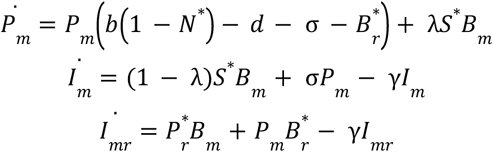

where *B*_*m*_ = β_*m*_ *I*_*m*_ + β_*mr*_ *I*_*mr*_.

The condition for mutant invasion, given by the condition for a positive real part of the dominant eigenvalue of the Jacobian obtained from the mutant population dynamics, is

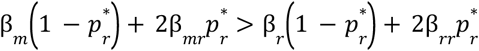

where 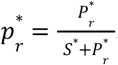 For every (α_*r*_, α_*m*_) pair, we used the within-host dynamics to compute the transmission rates β_*r*_, β_*m*_, β_*rr*_ and β_*mr*_, and numerically integrated the resident population dynamics for every value of α_*r*_ to obtain 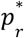. Using these values, we constructed the pairwise-invasibility plot shown in Figure 2 based on whether the mutant invasion condition is satisfied or not.

### Lattice model

#### Initialization

We constructed a 200×200 square lattice with wrapped boundaries. Initially, every lattice point is occupied by a susceptible host. Half of them carry a provirus and are assigned the state *P* and the other half are assigned the state *S*. The α value of each provirus is a random number drawn between zero and one.

During every timestep of the simulation, all the lattice points are updated once synchronously based on the following Next State Function.

#### Next State Function

1. If a lattice site is empty, we pick another lattice site randomly from the full grid with probability *G*_*R*_ or from the Moore neighbourhood with probability 1 − *G*_*R*_. If the chosen lattice site is occupied by a host, it reproduces into the empty site with probability *b*. If the host carries a provirus, it is always inherited by the offspring.
2. If the lattice site is occupied by a susceptible host
  - It can die with probability *d* and leave an empty site.
  - It can be infected by another host. We pick a random site from the full grid with probability *G*_*T*_, or from the Moore neighbourhood with probability 1 − *G*_*T*_. The focal host is infected if the chosen host is in state I and with the horizontal transmission probability of the virus(es). If the focal host is in state *S* and it is infected, it switches to state *P* with probability λ and the acquired virus is converted into a prophage. Otherwise, with probability 1 − λ, the host switches to state *I* where the acquired virus is in a replicating state. If it is chosen to infect a susceptible host in the following timesteps, it can transmit this virus (of trait value α_*i*_) with horizontal transmission probability β_*i*_. If the focal host is in state *P*, upon successful infection, it always switches to state *I* with two replicating viruses: its activated provirus and the acquired virus. If it is chosen to infect a susceptible host in the following timesteps, it can transmit either one of the two viruses (of trait values α_*i*_ and α_*j*_), with horizontal transmission probabilities β_*ij*_ and β_*ji*_ respectively.
  - If the focal host is in state *P*, then its provirus is activated with probability σ upon which the host switches to state *I*.
3. If the lattice site has a host in state *I*, it dies with probability γ.

#### Demonstration experiment

To demonstrate the selection for low-α provirus in Figure 4, we first initialised the lattice with hosts in state *S*. Then we converted a circular patch of host in the centre of the grid to state *I* with a replicating virus of α = 0. 6. At equal distances from the centre of the grid, we converted six circular patches of hosts into state *P*. The proviruses in these six patches were given six different α values (0.0, 0.2, 0.4, 0.6, 0.8 and 1.0). To focus on the survival of the provirus as a function of its α value, we set mutation, integration and provirus activation rates to zero.

### Virus factory model

In the virus factory model, the within-host virus replication dynamics and the host-level dynamics are interlocked by updating them together at every timestep. In addition, we explicitly modelled the extracellular virus dynamics, thereby relaxing the quasi steady state assumption. To achieve this, we consider an additional lattice that is overlaid on the host lattice. On this extracellular virus lattice, every lattice point can hold multiple individual virus particles which make up the local neighbourhood of the host occupying the overlying lattice point on the host lattice. For a susceptible host, each of the virus particles in its local neighbourhood can enter with an entry probability β. All of the viruses that entered the host can integrate and form proviruses with a fixed integration probability λ. If they do not integrate, the viruses will set up a virus factory. If a virus factory is set up, the host stops reproducing, loses its susceptibility, and the viruses that entered the host begin replicating probabilistically. The viruses within a factory maintain discrete population sizes and follow the same public goods dynamics, i.e. the replication probability is proportional to the average α value within the factory and a cost factor scaled by the α value of the virus. Whenever a virus replicates, the new copy has a certain probability of acquiring a mutation in its α value drawn from a Gaussian distribution around the parent α value, but capped at zero and one. At every time step, each provirus of the cell has a certain excision probability *e* to enter the virus factory where it will replicate alongside the other viruses. After a fixed replication period *T*_*L*_, the host dies and releases all the virus particles within the factory into its local neighbourhood. On the extracellular virus lattice, each particle has a diffusion probability of moving to a neighbouring lattice point and it is lost from the system with a certain decay probability. A susceptible host, if it carries proviruses, can also spontaneously set up a virus factory that is initialised by all of its proviruses. See Supplementary Information S6 for a complete description of the model.

The individual based models were implemented using Cacatoo (van Dijk, 2022). Codes are available on github.com/sultannazir/proviral_defense_system. An interactive version of the virus factory model is available on sultannazir.github.io/proviral_defense_system.

#### Parameter sweep

We explored the virus factory model (with many more interesting parameters) using Latin Hypercube Sampling. We generated 10,000 random parameter combinations (using SciPy (Virtanen et al., 2020)) within reasonable bounds (see Supplementary Information S1), and classified whether or not defense systems emerged. Each parameter combination was simulated once, with its own random seed, for 200,000 timesteps.

#### Characterising the emergence of a defense system

For every simulation run, we classified the outcome as ‘selection for defense’ or ‘defense evolved’, in Figure 3(C) and Figure 5(C) respectively, if over 20% of all viruses are proviruses that have an α value less than 0.1.

## ACKNOWLEDGEMENTS

This collaborative project was made possible by the Swaantje Mondt travel fund for complexity research awarded by the Centre for Complex Systems Studies (CCSS) at Utrecht University.

## GEN-AI STATEMENT

We used Claude Opus 5 (Anthropic) to port the virus factory model to a lower-level implementation, which made the large-scale parameter sweeps computationally feasible. First, the ported code was validated against our original prototype, reproducing its output bit-for-bit. The authors take full responsibility for this code, the results obtained with it, and the further content of this manuscript.

## AUTHOR CONTRIBUTIONS

S.A.N. designed the model with input from B.A. and B.v.D., and programmed the model. S.A.N. and B.A. derived the mathematical results. S.A.N. and B.v.D. simulated the model and analysed data. S.A.N., B.A., A.L. and B.v.D. interpreted the results and wrote the paper.

## SUPPLEMENTARY INFORMATION

### S1: Table of parameters

**S7: Virus factory model parameter sweep**
| Model parameter | Meaning | Adaptive dynamics | Lattice model |
| --- | --- | --- | --- |
| $b$ | Intrinsic host birth rate/probability | 0.05 | 0.01 |
| $d$ | Intrinsic host death rate/probability | 0.01 | 0.002 |
| $\beta_0$ | Infection term coefficient | 1.0 | 1.0 |
| $\lambda$ | Virus integration rate/probability | 0.05 | 0.01 |
| $\sigma$ | Provirus spontaneous activation rate/probability | 0.0005 | 0.0001 |
| $\gamma$ | Host lysis rate/probability | 0.05 | 0.01 |
| $r$ | Maximum virus replication rate | 0.25 | 0.25 |
| $c$ | Cost of cooperation | 0.5 to 1.0 | 0.5 to 1.0 |
| $T_L$ | Virus replication period | 20 | 20 |
| $\theta$ | Time unit conversion coefficient | 1.0 | 0.2 |
| $G_R$ | Probability of global host reproduction | N/A | 0.0 to 1.0 |
| $G_T$ | Probability of global virus transmission | N/A | 0.0 to 1.0 |

| Virus factory model parameter | Default value | Sweep range |
| --- | --- | --- |
| Host birth probability, $b$ | 0.05 | - |
| Susceptible host death probability, $d$ | 0.01 | - |
| Free virus entry probability, $\beta$ | 0.5 | 0.1 to 1 |
| Integration probability, $\lambda$ | 0.1 | 0.001 to 0.1 |
| Probability of spontaneous provirus activation, $\sigma$ | 0.0001 | $10^{-5}$ to $10^{-3}$ |
| Probability of a provirus joining a virus factory, $e$ | 0.1 | 0.01 to 1 |
| Maximum probability of virus replication within factory, $r$ | 0.25 | - |
| Cost of cooperation, $c$ | 0.5 | - |
| Virus replication period, $T_L$ | 20 | - |
| Probability of virus mutation, $\mu_\alpha$ | 0.001 | - |
| Standard deviation of mutation, $\mu_{SD}$ | 0.1 | - |
| Probability of provirus loss during host reproduction, $l$ | 0.001 | - |
| Virion diffusion probability (if $<1$) or Number of diffusion events per virion per timestep (if $\geq 1$), $D_v$ | 0.5 | 0.1 to 100 |
| Extracellular virus decay probability, $\delta_v$ | 0.05 | 0.01 to 0.4 |

### S2: Derivation of optimum α using adaptive dynamics

Following the ODEs for the mutant dynamics given in the Methods section, the Jacobian is

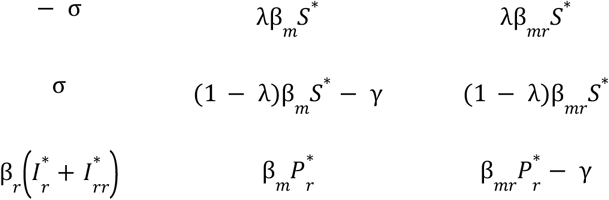

Solving for the three eigenvalues, we find that one of the eigenvalues is

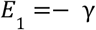

The other two eigenvalues are given by the solution to

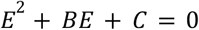

where 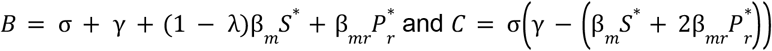

Because *B* > 0, using the quadratic formula, the condition for mutant invasion (a positive leading eigenvalue) is reduced to *C* < 0.

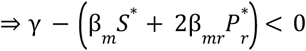

From the steady state condition of resident dynamics, we get 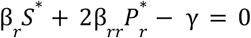. Substituting and dividing by 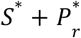, the condition of mutant invasion is

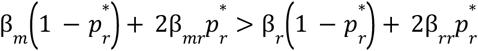

For our model, β_*r*_ = β_*rr*_. Therefore, the condition becomes

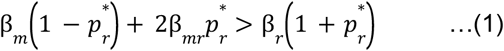

Assuming quasi steady state of viral infection dynamics, the horizontal transmission rate is simply proportional to the burst size of the viral lineage. Let *R* α_*x*_, α_*y*_ be the burst size of a α_*x*_ valued virus when coinfecting with a α_*y*_ valued virus. Note that for our framework of within-host dynamics, the burst size of a singly infecting lineage is equal to its burst size when coinfecting with a virus of the same trait value.

Therefore, our condition for mutant invasion from above can be rewritten as

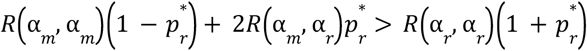

Let us define the invasion fitness of the mutant as

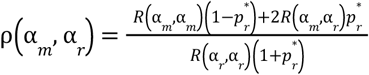

The condition for positive selection on α is 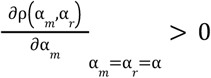

Evaluating (*R* α_*m*_, α_*r*_) around α_*m*_ *≃* α_*r*_ and substituting on our within-host framework gives us

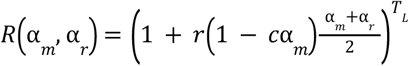

Substituting and simplifying, the condition for positive selection on α reduces to

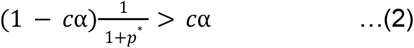

which takes the form of Hamilton’s rule (Hamilton, 1964) where 1 − *c*α is the marginal benefit of cooperation, *c*α is the marginal cost and 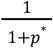 is the average relatedness.

At the optimum trait value, the selection gradient is zero. Hence we get,

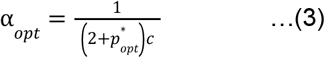

Since selection is positive below this value and negative above value, we have converging evolution towards this optimum.

#### Derivation with lineage-weighted averages

For the analysis in this study, we defined the state variable 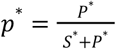 which is the equilibrium fraction of susceptible hosts that carry a provirus. However, we could instead consider a *lineage-weighted* fraction which accounts for the fact that hosts with proviruses carry an additional viral lineage, and define 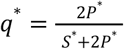.if *p* is thought of as the chance of coinfection across hosts, *q* ^*\**^ is the chance for coinfection per viral lineage.

With this modification, the invasion condition (1) becomes

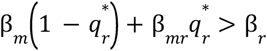

This means that a mutant virus invades if its *lineage-weighted* average rate of horizontal transmission is higher than the resident virus.

The condition for positive selection on α based on Hamilton’s rule (2) becomes

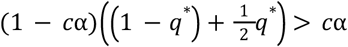

The expression of relatedness here is 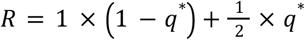. Given that the relatedness within single infections and coinfections are 1 and 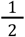 respectively, and *q*^*^ is the per-lineage chance for coinfection, we can clearly see here that the relatedness measure we obtained is the average relatedness across viral lineages at equilibrium.

Accordingly, the expression for the optimum trait value (3) becomes

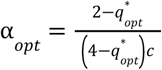

### S3: *R*_*0*_ derivation

Consider a host population without any virus whose dynamics is described by

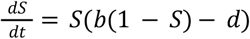

which goes to a steady state value of 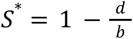.

The invasion dynamics of a virus is described by

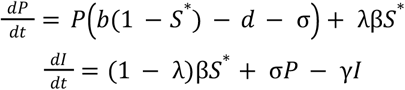

Where we can ignore the possibility of coinfections because the virus is rare.

The Next Generation Matrix of the virus is given by

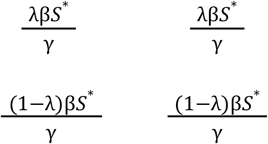

whose eigenvalues are 0 and 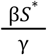. Therefore,

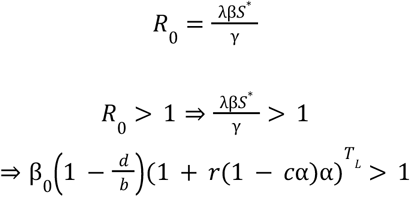

Simplifying, we get lower and upper bounds on the value of α given by

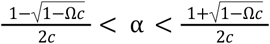

where 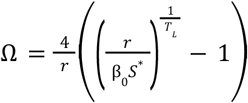.

### S4: Lattice model time series

Figure S1 shows the time series of the frequency distribution of α for different values of the cost of cooperation *c*. We observed that there is a stronger selection for the defense system for smaller values of *c*. This is because lower cost of public good production also means a higher maximal replication rate for the cooperating viruses. Therefore, we have larger burst sizes of the parasites and a stronger selection pressure for defense.

**Figure S1:**
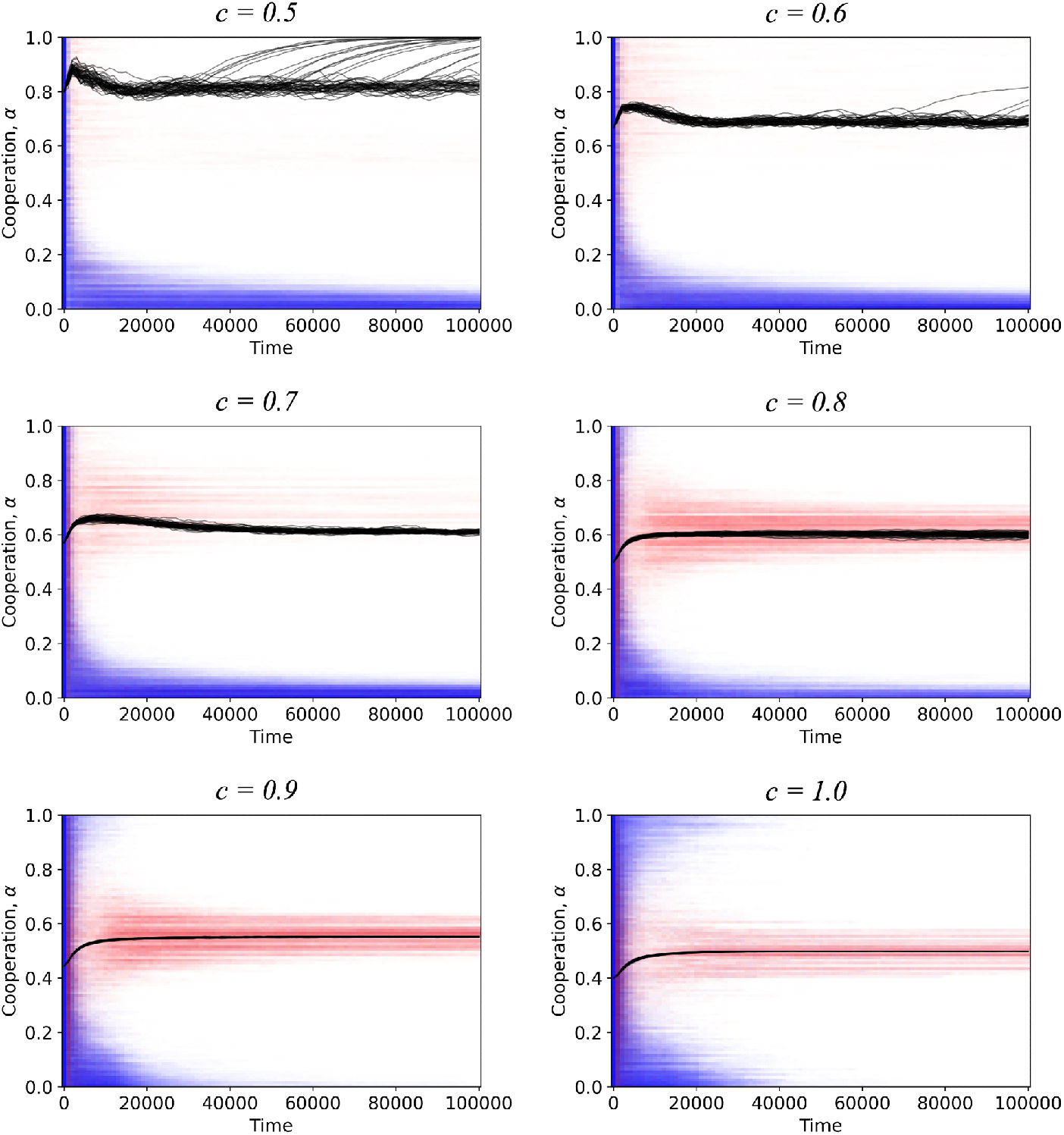
Lattice model initialised at a uniform distribution of α value and simulated without mutations. Overlaid α distributions of viruses from 50 simulation replicates, for 6 different values of cost of cooperation *c*. The frequency distributions of free viruses and proviruses are plotted in red and blue respectively, which appear purple when they overlap. Both the free virus and provirus distributions, at every timepoint, are normalised by the peak of their cumulative distribution, i.e. the total virus frequency distribution. The black lines denote the predicted optimum α given by 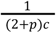 where *p* s the fraction of susceptible hosts carrying a provirus at that timepoint. The distributions shown in Figure 3(B) are obtained from the averages over timepoints 90,000 to 100,000 of these simulations. *G*_*R*_ = 0, *G*_*T*_ = 0

### S5: Lattice model with mutations

Figure S2 shows that the defense systems evolved in our lattice model *de novo* but only when the mutation rate (µ_α_) and mutational step size (µ_*SD*_) are high. For cost of cooperation *c* = 0. 5, we see that defense systems evolve consistently across all replicates for high mutation rate and high mutation step size. This ensures that there is plenty of diversity in the virus population for selection to act on a lineage of proviral cheats. However, for the same mutation parameters, defense did not evolve for *c* = 0. 8 even when the simulations were able to generate traces of proviral cheats. This may be due to some frequency dependence and/or the weaker strength of selection associated with higher values of *c*.

**Figure S2:**
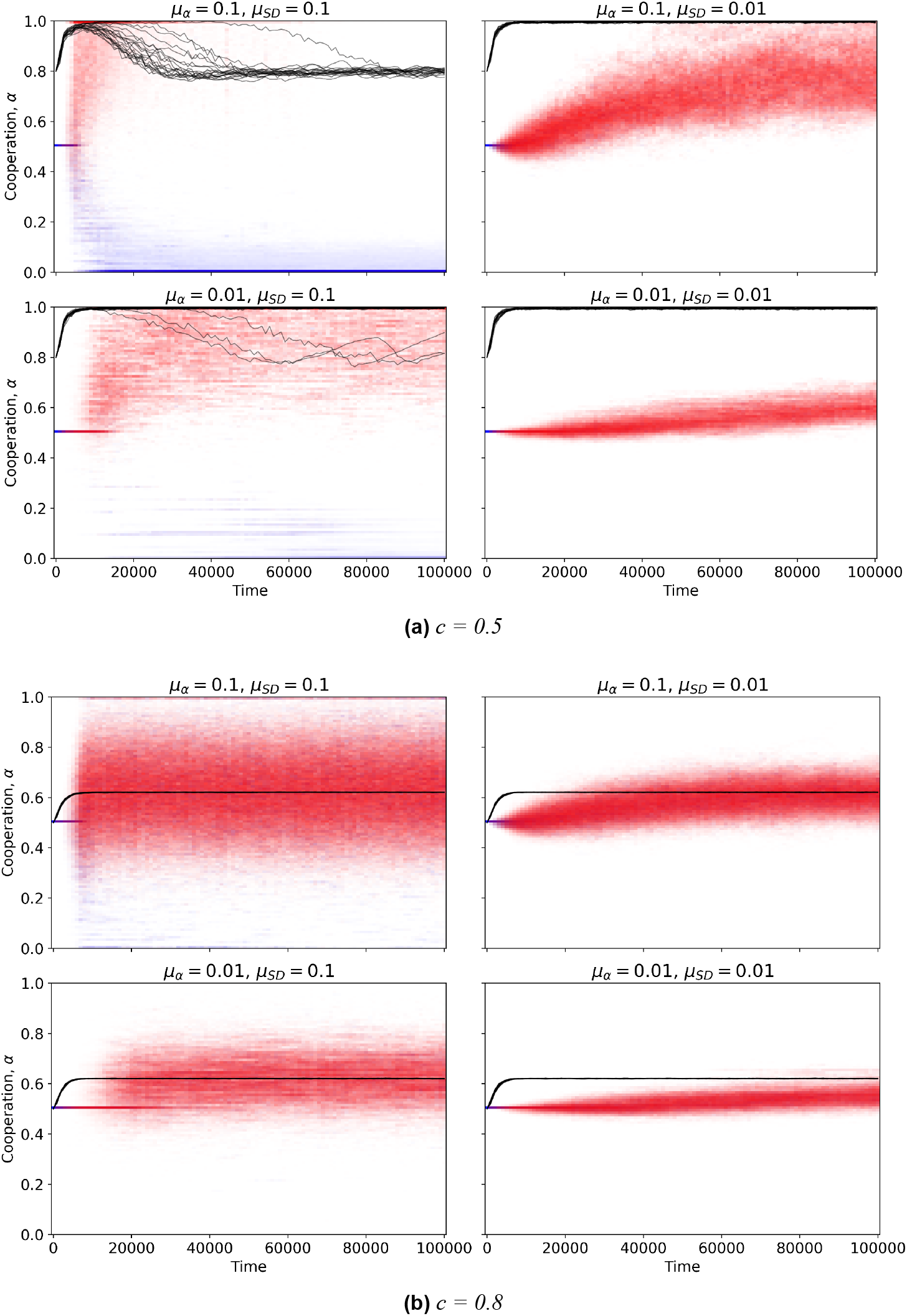
Lattice model initialised at a single α value and simulated with mutations. Overlaid α distributions of viruses as described in Figure S1, but with 20 replicates of simulation runs testing for *de novo* evolution through mutations. The runs are initialised with a virus of α = 0. 5 and different values of the mutation parameters µ_α_ (probability of mutation) and µ_*SD*_ (standard deviation of mutation step size). *G*_*R*_ = 0, *G*_*T*_ = 0

### S6: ODD-protocol of virus factory model

Because the virus factory model is a relatively complex, individual-based model, we describe this using the ODD (Overview, Design concepts, Details) protocol (Grimm et al., 2020) to improve reproducibility.

1. **Purpose** With the virus factory model, we studied the *de novo* evolution of proviral cheats that act as a host defence system, primarily with more biological realism than the lattice model where all within-host replication was implicit. Per the lattice model, spatial structure remained the focal explanatory variable, but other elements related to diffusion and timing were further explored.
2. **Entities, state variables, and scales** Hosts are cells on a square lattice (200 × 200) with periodic boundaries and a Moore neighborhood (8 surrounding lattice sites). Each cell has a state that is either {empty, susceptible, infected} and a clock counting timesteps since factory formation. Virus elements are individuals carrying one evolvable trait, α ∈ [0, 1]. Each lattice site holds three independent lists:
  - Integrated proviruses (in every lattice site with a host)
  - Factory viruses, those replicating inside an active factory (only in sites with an infected host)
  - Extracellular free virions at that location Time is discrete and has arbitrary units. Runs typically lasted 200,000 timesteps.
3. **Process overview and scheduling** Each timestep, in order:
  1. (*mixed treatment only*) the host grid is randomly permuted.
  2. Host update. One sweep over all cells, reading host states from a copy of the previous state. (synchronous update, as internal hosts dynamics does not influence other hosts)
  3. Extracellular update. Each free virion decays or diffuses. (asynchronous update, needed to ensure mass conservation with diffusion modifying other lattice sites) Within the host update, behaviour depends on cell state:
  - **Susceptible:** dies with probability *d*. Otherwise each co-located free virion enters with probability β; if any entered, a single draw decides whether they form a factory (cell → infected, clock = 1) or integrate as proviruses. If the cell is still susceptible and carries proviruses, spontaneous induction fires with probability σ, moving every provirus into a new factory.
  - **Infected:** each provirus excises and joins the factory with probability *e*. Every factory element then replicates (below). clock increments; when clock hits the latency timer, the cell bursts: all factory contents are released as free virions at that lattice site and the host is removed (lattice site ‘empty’)
  - **Empty:** one random neighbor is drawn from 8 surrounding lattice sites; if susceptible, it reproduces into the cell with probability *b*. The daughter inherits each parental provirus independently with probability (1 − *l*) where *l* is the probability of provirus loss.
4. **Design concepts** Basic principles - public-goods cooperation and tragedy of the commons for viruses within a host, spatial structure as the mechanism to sustain cooperative traits Emergence - the alpha-distribution emerges through evolution within the viral population, and emergent spatial patterns uphold a degree of relatedness amongst hosts Adaptation - the α values of viruses undergo mutations and selection Sensing - strictly local: hosts only interact with viruses at the same lattice site, and only compete with nearby individuals for empty spots Stochasticity - mutation, reproduction, and infection are all events based on probabilities, done with a seeded random number generator Collectives - viruses in factories form a collective, jointly influencing one another’s replication rates. In addition, spatial structure forms collectives of related hosts (and their proviruses) Observation - Every 100 steps, host counts (S, I), 100-bin α histograms of the provirus pool and of the free-virion pool separately, and the distribution of provirus copy number per host. For the parameter sweeps, runs are additionally classified into host extinct, virus extinct, no defense, or defense evolved.
5. **Initialisation** All cells are initially susceptible. Each cell independently receives one provirus with probability 0.5, and each lattice site receives one free virion with probability 0.01; every initial element has α_0_ = 0. 5. All clocks start at one.
6. **Input data** No external forcing data was used
7. **Submodels**
  7.1 **Replication in the factory**: with 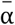 the factory mean, element *i* produces one offspring with probability 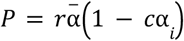. The benefit term depends on the *group*, the cost term on the *individual*, so low-α viruses out-replicate their factory-mates while depressing 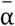 for everyone.
  7.2 **Mutation:** on every copy (factory replication and provirus inheritance) α mutates with probability µ_α_ by a Gaussian step of s.d. step µ_*SD*_, truncated to [0, 1].
  7.3 **Diffusion and decay:** each timestep each free virion decays with probability δ_*V*_; survivors may move to one adjacent lattice site (including diagonals) with probability *D*_*V*_. If the value of *D*_*V*_ is above one, we implement *D*_*V*_ many moves of each virion to a randomly drawn adjacent site at every move, followed by stochastic rounding. Under the mixed treatment virions are instead moved to a random lattice site.
  7.4 **Observation:** population statistics are collected (see part 4)
8. **Implementation** The model was developed and prototyped with the Cacatoo framework (van Dijk, 2022). However, all sweeps reported in the main text were produced by a Python/Numba reimplementation, written with assistance of Anthropic Claude, purely as an optimisation. The model logic was confirmed to track our prototype’s by byte-comparison. Codes are available here: https://github.com/sultannazir/proviral_defense_system

### S7: Virus factory model parameter sweep

**Figure S3:**
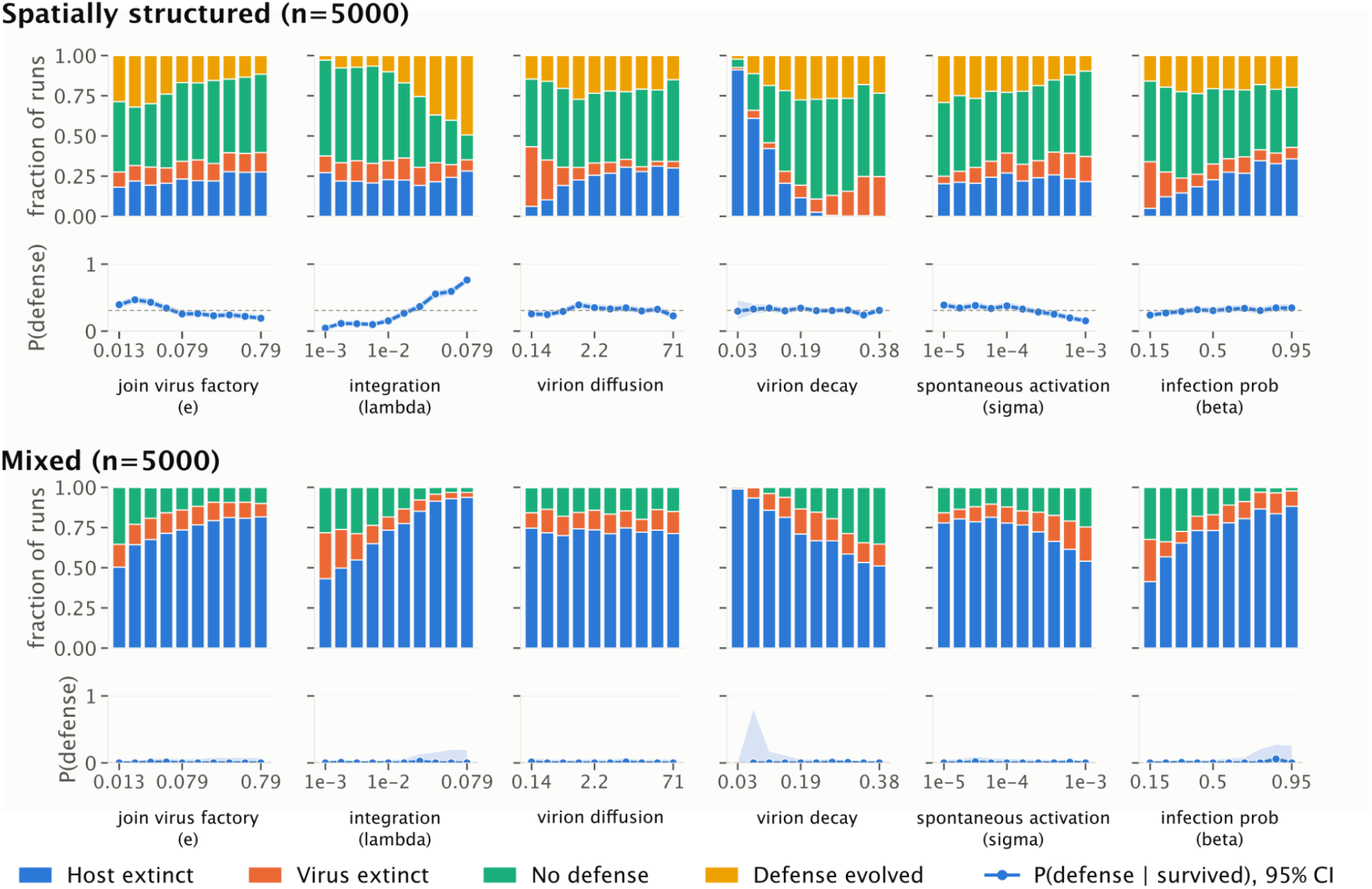
Virus factory model sensitivity analysis. Projection of outcomes of simulation runs from a Latin hypercube sampling of model parameters onto each of the parameters. Six parameters were varied (see Table S1 in the main text): the probability of a provirus joining a virus factory (0.01 to 1, log-uniform), the integration probability (0.001 to 0.1, log-uniform), the extracellular virus diffusion probability (0.1 to 100, log-uniform), the extracellular virus decay probability (0.01 to 0.4, uniform), the probability of spontaneous provirus activation (10^−5^ to 10^−3^, log-uniform), and the free virus entry probability (0.1 to 1, uniform). All other parameters were held at the default values listed in Table S1. The top row shows the parameter sweep in spatially structured host populations, and the bottom row the same sweep for a well-mixed grid where the hosts positions are shuffled at every timepoint of the simulation. We observe that spatial structure of the host is indeed necessary for the evolution of the defense system. Note that although *e* and *D*_*v*_ (‘virion diffusion’ here) do not appear to have a strong effect on defense evolution here, this a binning-artefact due to their combined effect (shown in Figure 5(C) in the main text) cancelling each other out.

